# Daytime eyeshine contributes to pupil camouflage in a cryptobenthic marine fish

**DOI:** 10.1101/177519

**Authors:** Matteo Santon, Pierre-Paul Bitton, Ulrike K. Harant, Nico K. Michiels

## Abstract

Ocular reflectors enhance eye sensitivity in dim light, but can produce reflected eyeshine when illuminated. Most teleost fish occlude their reflectors during the day. The opposite is observed in cryptic sit-and-wait predators such as scorpionfish and toadfish, where reflectors are occluded at night and exposed during the day. This results in daytime eyeshine, proposed to enhance pupil camouflage by reducing the contrast between the otherwise black pupil and the surrounding tissue. In this study, we test this hypothesis in the scorpionfish *Scorpaena porcus* and show that eyeshine is the result of two mechanisms: the previously described *Stratum Argenteum Reflected* (SAR) eyeshine, and *Pigment Epithelium Transmitted* (PET) eyeshine, a newly described mechanism for this species. We confirm that the ocular reflector is exposed only when the eye is light-adapted, and present field measurements to show that eyeshine reduces pupil contrast against the iris. We then estimate the relative contribution of SAR and PET eyeshine to pupil brightness. Visual models for different light scenarios in the field show that daytime eyeshine enhances pupil camouflage from the perspective of a prey fish. We propose that the reversed occlusion mechanism of some cryptobenthic predators has evolved as a compromise between camouflage and vision.

## Introduction

An ocular reflector behind the retina is a common feature of vertebrate eyes. Their presence and diversity across taxa is linked with increased visual sensitivity under dim light^1^. This is achieved by reflecting light not captured by the retina during its first pass back through the photoreceptors, allowing for an increased photon catch^2–4^. However, ocular reflectors come with two disadvantages. First, visual acuity might be reduced by backscatter in bright environments^5–7^. Secondly, an animal may become more conspicuous because of eyeshine caused by reflection of light out of the pupil^7^. In most teleosts and a few reptiles, these side effects are minimized by occlusion mechanisms that cover the reflector in the light-adapted eye with black melanin pigmentation^1^. Hence, strong eyeshine can only be induced when the eye is dark-adapted; it is weak or absent when the eye is light-adapted.

In contrast to this general pattern, a few families of highly cryptic fishes such as toadfishes (Batrachoididae) and scorpionfishes (Scorpaenidae) feature strong eyeshine when the eye is light-adapted, but not when dark-adapted^6,8,9^. In these species, the reflector is a *stratum argenteum* located in the outer part of the choroid^1,6,10^, rather than the common *tapetum lucidum*. This *stratum argenteum* is a bi-laminate reflective structure with an inner layer of pentameric uric acid crystals and an outer layer of yellow birefringent granules of unknown material^9,11^. In the light-adapted state of the eye, the melanosomes of the retinal pigment epithelium enter the cell processes between the cones, clearing the way for light that passed through the receptor layer to also penetrate the translucent choroid^1,6^ and reach the *stratum argenteum*. This light is reflected out of the pupil, generating a type of eyeshine termed *Stratum Argenteum Reflected* (SAR) eyeshine^12^. Another consequence of choroid translucency is that down-welling light that penetrates the dorsal part of the eye can be transmitted through the sclera and choroid and out through the pupil. This can result in a second type of eyeshine described as *Pigment Epithelium Transmitted* (PET) eyeshine^12^. While the translucency of the sclera and choroid has been already documented for these fishes^9,13^, pupil eyeshine has always been assumed to be the exclusive result of SAR eyeshine. The possibility that PET eyeshine also contributes to total eyeshine has not been considered. Both eyeshine types might explain why scorpionfishes, toadfishes and some stonefishes have bright pupils when exposed to ambient light^13^. The possible function of this counter-intuitive occlusion mechanism, however, is yet unclear. Here, we investigate the hypothesis that it conceals the pupil by allowing daytime eyeshine and thus reducing contrast with the surrounding tissue^9^(video S1).

The well-defined, often circular, dark pupil that stands out against the body in most animals makes the vertebrate eye difficult to hide^14^. Eyes are indeed considered key features for face recognition of predators, prey or conspecifics^15–21^. Consequently, mechanisms for pupil camouflage are widespread. Pupillary closure (e.g.elasmobranchs) reduces pupil size and shape in response to fluctuations in ambient light, but also minimizes pupil conspicuousness^14^. Lidless species, such as fishes and snakes, often feature an eye mask that embeds the pupil in a dark skin pattern (e.g. vertical stripes in lionfishes)^14^. In some fish species, iridescent corneal reflectors have been proposed to reduce an eye’s detectability^22^. Skin flaps (e.g. flatheads), where the pupil is partly covered by an irregular extension of the cryptic iris, are another adaptation to reduce conspicuousness^14^. Some fishes simply have very small pupils for their body size (e.g. frogfishes), which may be a strategy to minimize eye detection at the cost of visual acuity. Thus far, the use of eyeshine for enhancing daytime eye camouflage has only been proposed for invertebrates: pelagic stomatopod larvae reduce the conspicuousness of their dark retinas by eyeshine that matches the light field of the background^23^. Scorpionfishes, toadfishes and stonefishes are sit-and-wait predators that strongly rely on crypsis. Since featuring a dark pupil could disrupt their camouflage, diurnal pupil eyeshine might help to hide their eyes.

After confirming that our model species, the black scorpionfish *Scorpaena porcus*, features a reversed occlusion pattern of the *stratum argenteum*, we describe pupil radiance in relation to the iris in the field using spectroradiometry. We then quantified the two different eyeshine mechanisms in the laboratory. These measurements were further used to model pupil contrast in relation to the iris and skin from the perspective of a prey fish under three light scenarios. This allows us to assess to what extent eyeshine could enhance pupil camouflage.

## Methods

### Model species

The black scorpionfish *Scorpaena porcus* is common in coastal marine hard-substrate and seagrass habitats in the eastern Atlantic Ocean and Mediterranean Sea^24^. It is a generalist sit-and-wait predator that relies on crypsis to catch naïve prey. Like most scorpionfish, it features prominent eyes with a translucent retinal pigment epithelium resulting in daytime eyeshine (Fig. 1)^6,12^. We caught 15 individuals in Calvi (Corsica, France) between 5 and 20 m depth under the general permit of STARESO (Station de Recherches Sous Marines et Océanographiques). At STARESO, fish were kept in two 300 L tanks with a continuous fresh seawater flow. For field measurements, three individuals were used and subsequently set free. The remaining twelve were transported to the University of Tübingen (Germany) in individual plastic canisters filled with 1.5 L seawater and oxygen enriched air. In Tübingen, fish were held individually in 160 L tanks (20°C, salinity 35 ppt, pH 8.2, 12h light/dark cycle, fed once every two days). Animal husbandry was carried out in accordance with German animal welfare legislation. Because the individuals were not experimentally manipulated, a formal permit was not required for this study (confirmed following conversations with Annette Denzinger, Animal Care Officer at the Biology Department of the University of Tübingen).

**Figure 1.**
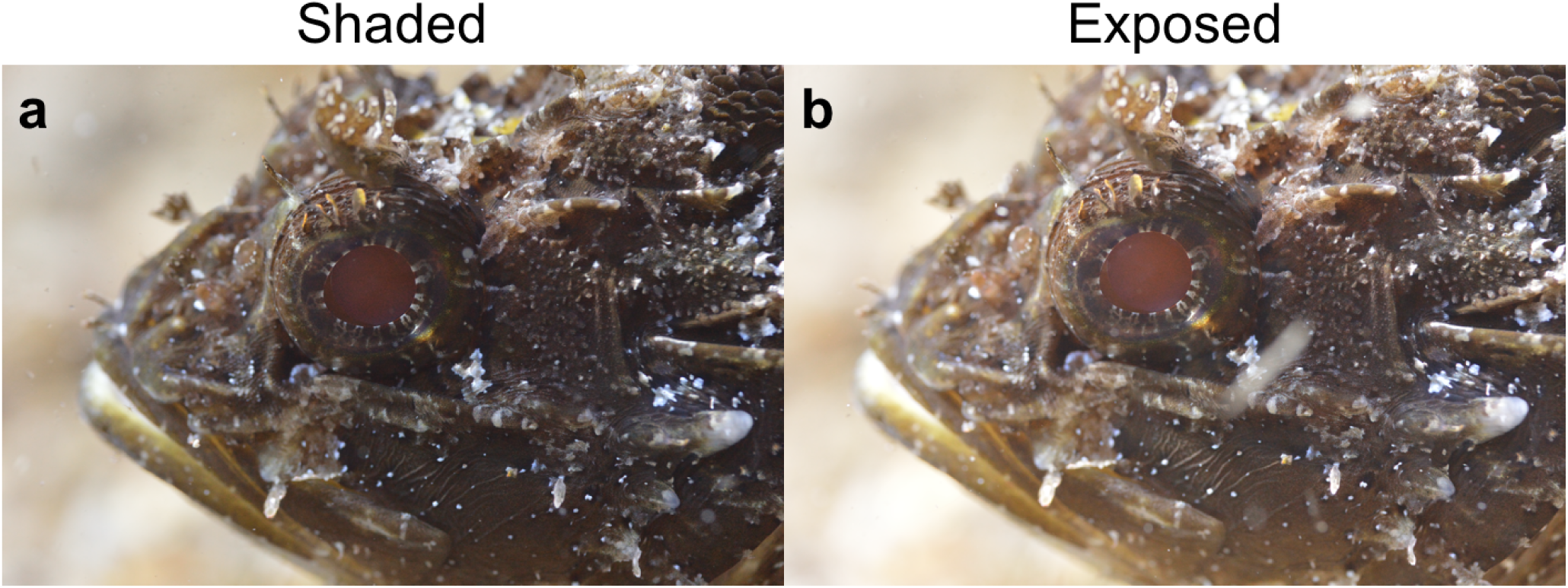
*S. porcus* under shaded and exposed conditions in the field. When the fish is shaded (a) or exposed (b) eyeshine seems to reduce pupil contrast against the iris. The overall brightness of the pictures is similar despite shading in A because of automatic exposure, illustrating the effect of shading on eyeshine relative to the nearby tissue.

### Spectroradiometry

#### Occlusion of the stratum argenteum during dark-adaptation (laboratory)

To confirm that the *stratum argenteum* is occluding while the eye is dark-adapting, an individual was placed in a 12 L tank (L × W × H: 15 × 15 × 20 cm) positioned on a cooling plate and equipped with an aeration stone. Before measuring, we light-adapted the fish for 2 h using a sun-simulating Plasma-i AS1300 Light Engine (Plasma International, Mühlheim am Main, Germany) pointed upwards to a diffuse reflector (#273 soft silver reflector, LEE Filters, Andover, England) attached to the ceiling to illuminate the whole room. Three polytetrafluoroethylene (PTFE) white reflectance standards (Lake Photonics, Uhldingen-Mühlhofe, Germany) were positioned sideways in the tank for ambient light measurements. We measured a total of three individuals.

Radiance measurements of the left eye pupil were obtained using a calibrated SpectraScan PR 740 spectroradiometer (Photo Research, NY, USA). This device uses Pritchard optics and measures the absolute spectral radiance of an area with known solid angle. The lens of the spectroradiometer was aligned orthogonally to the pupil of the fish, and the tank was tilted at an angle of 5° to reduce external reflection. We used a cold light source KL2500 LCD (Schott, Mainz, Germany) equipped with a blue filter (insert filter 258302, Schott, Mainz, Germany) to coaxially illuminate the pupil of the fish. The light was led through liquid light guides (LLG 380, Lumatec, Deisenhofen, Germany) to a mechanical shutter and then on to a 90° elbow-shaped glass-fibre light guide (Heine Optotechnik, Herrsching, Germany) with an exit diameter of 3 mm. The light exit was aligned coaxially with the spectroradiometer’s optical axis, 15 cm in front of the lens. The distance between the lens and the fish was fixed at ∼50 cm. Slight distance adjustments were necessary to assure complete coverage of the pupil with the cross-section of the measuring area. Since scorpionfish tend to sit passively on the substrate, there was no need for the use of anaesthesia, which is known to affect fish pigmentation^25^.

Measurements started immediately after turning off the plasma source. The room was kept completely dark except for brief moments (< 10 s) where the shutter of the coaxial source was opened for measuring, approx. once every 10 min for 2 h. The radiance of the PTFE white standard best aligned with the fish’s pupil was measured orthogonal to its surface at the beginning and the end of the experiment. SAR eyeshine reflectance was calculated as the photon radiance of the pupil normalized (i.e. divided) by the average radiance of the PTFE white standard. Because it was exclusively generated by a coaxial light source in an otherwise dark environment, a condition that would be uncommon in the field and that was explicitly staged to strictly describe the reflective properties of the *stratum argenteum*, we subsequently refer to this measurement as “narrow-sense” SAR eyeshine.

We analysed the data using a log-linear model with total reflectance (reflectance integrated over the wavelength range from 380 to 780 nm) as response variable, and dark-adaptation time and individual ID as predictors using R v3.2.0^26^. Model assumptions were verified by plotting residuals versus fitted values and versus each covariate in the model. Significance of predictors was tested by using the 95% Credible Interval (CrI), a Bayesian analogue of the confidence interval^27^. For the response variable time, the factor individual ID, and their interaction, we computed the model estimates from the back-transformed effect sizes. The associated 95% CrI were then obtained from 10000 simulations of the mean and variance of each estimate, using the sim function of the R package arm, with non-informative priors^28^. If the CrI of one individual did not overlap with the mean of another, we concluded that their intercepts were significantly different. We considered time and its interaction with ID to be significant if the CrI values of the log-linear regression coefficient did not include zero.

#### Contrast of natural and model pupil against iris (field)

To confirm that *S. porcus* produces enough daytime eyeshine to reduce pupil contrast against the iris in the field, we measured pupil and iris radiance of three fish in situ in STARESO. Each fish was measured in a transparent plastic terrarium (L × W × H: 15 × 20 × 15 cm) placed underwater. The side of the container through which measurements were taken consisted of Evotron optically neutral Plexiglas^®^ (Evonik Performance Materials, Essen, Germany). A black PVC sheet was added as a background to encourage the scorpionfish to face the spectroradiometer. Holes in the sides ensured water circulation. As an optional shade, we used a black plastic slate. All measurements were taken using the SpectraScan PR 740 in a custom-built underwater housing (BS Kinetics, Achern, Germany) with the spectroradiometer facing south and the fish north, at noon on a clear, sunny day. The focal distance of the instrument was fixed at 80 cm. The external dimensions of the underwater housing are 35 × 25 × 25 cm (L × W × H), excluding the length of the port (10 cm). The port is located at the upper edge of the housing. It is hard to evaluate the influence of the housing on the light field, but even if a partial obstruction would occur this would hardly influence our estimates since every radiance measurement was obtained under comparable geometries.

We measured pupil and iris radiance (top, bottom, left and right) in the same three fish at 7 m and 15 m depth. Under shaded conditions, pupil radiance involves SAR reflected eyeshine only, as PET transmitted eyeshine is prevented. Under exposed conditions, both mechanisms contribute to eyeshine. We also estimated the radiance of the skin patches surrounding the iris (top, bottom, right and left of the eye) for visual modelling (see below). As a reference, we measured three PTFE white standards, one facing upwards measured from a 45° angle as a proxy for down-welling light. Two more were facing the observer, one of them exposed and the other shaded, and were measured frontally as a proxy for side-welling light. Finally, we also measured a model pupil (Fig. S1). The latter consisted of a black, hollowed-out PVC block (L × W× H: 6 × 3× 3 cm) with a pupil-like opening at the front and internally filled with black cloth to absorb as much light as possible. The resultant pupil is practically black, mimicking a pupil with no eyeshine. We cycled through all measurements two times, both under shaded and exposed conditions.

To compare the contrast of the natural and dark model pupil against the iris, we calculated the average photon radiance of all three structures for each wavelength and exposure. We then calculated the Michelson contrasts Cm as follows:

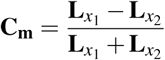

Where **L**_*x*1_ is the radiance of one of the two pupils, **L**_*x*2_ the radiance of the iris. There is no contrast when **C_m_ = 0**. When **C_m_** > **0**, **L**_*x*1_ is brighter than **L**_*x*2_, and vice versa for **C_m_** < **0**.

#### PET and narrow-sense SAR eyeshine in light-adapted fish (laboratory)

To describe the relative contributions of transmitted (PET) and narrow-sense reflected (SAR) eyeshine to pupil radiance in light-adapted scorpionfish, we measured 10 individuals in a similar setup to the one used to confirm occlusion during dark-adaptation. We used a down-welling tungsten source (650+ ARRI, Munich, Germany) attached to the ceiling pointing downward at the setup. The light reflected by the diffuse grey metal walls of the room generated the side-welling light. We assessed the contribution of PET eyeshine as the difference in pupil radiance between exposed and shaded conditions. PET eyeshine transmittance was subsequently obtained by dividing the difference between exposed and shaded pupil radiance by the radiance of the upward-facing, exposed PTFE white standard. To estimate the exclusive contribution of SAR eyeshine, we measured pupil radiance using the coaxial illumination system described earlier while only briefly turning off the main source (ARRI) to keep fish light-adapted. As a reference for down-welling light, we measured a PTFE standard facing upwards from an angle of 45°. Side-welling light was estimated by frontally measuring a PTFE white standard facing sideways. Narrow-sense SAR eyeshine reflectance was calculated as coaxially induced pupil radiance divided by the radiance of the coaxially illuminated PTFE white standard facing sideways.

#### Broad-sense SAR eyeshine, iris and dark model pupil reflectance for visual modelling

In the field, SAR eyeshine never occurs under the conditions used to strictly characterise the reflective properties of the *stratum argenteum* in the laboratory: light-adapted fish, sitting in the darkness, illuminated only by a coaxial light source. Hence, to properly reconstruct the contribution of SAR eyeshine assessed in the field, we also estimated “broad-sense” SAR eyeshine by measuring a fish and a PTFE white standard in the shade (PET eyeshine suppressed), without using a coaxial light source. In this setup, broad-sense SAR eyeshine is expressed in relation to the side-welling ambient light, as pupil radiance was assessed in the field. We measured nine fish under the down-welling ARRI light source and shaded them with a black PVC panel. As a proxy of the side-welling light field, we frontally measured a shaded PTFE white standard facing sideways in the tank. Broad-sense SAR eyeshine reflectance was calculated as the photon radiance of the pupil normalized by the radiance of the shaded PTFE white standard. We also measured the reflectance of the iris and the dark model pupil relative to a PTFE white standard in the same setup under exposed conditions for later visual modelling (see below).

To determine if our laboratory estimates of PET, broad sense SAR eyeshine and iris reflectance could be used to reliably predict field data, we used these measurements to ‘reverse engineer’ the pupil and iris radiance for three fish for which complete field data were available: one at 7 m and two at 15 m. We used side- and down-welling light measurements from the field as the illuminants. By combining them with the broad-sense SAR, PET eyeshine and iris relative radiances estimated in the laboratory, we then predicted pupil and iris radiance under shaded or exposed conditions for each of the three fish. We then assessed the match between the two curves by calculating the mean of the ratio between the logarithm on base 10 of the predicted and real radiances at each wavelength for iris and pupil of each fish under both exposures. Finally, we computed the averaged ratio for all three fish.

### Visual models to assess pupil camouflage as perceived by prey fish

To test if PET and SAR eyeshine enhance pupil camouflage, we determined how well the pupil matches the appearance of the iris as perceived by one of its prey species, the triplefin *Tripterygion delaisi*^29^ under three light scenarios. We calculated the chromatic contrasts between the pupil and the iris using the receptor-noise model^30^. We informed the model using species-specific visual system parameters including the visual sensitivity of the photoreceptors (SWS: 468 nm, MWS: 517 nm, LWS: 530 nm), the relative photoreceptor densities in the fovea of 1:4:4 (SWS : MWS : LWS), the ocular media transmittance, and setting the Weber fraction at 0.05^29–31^. The achromatic contrasts were calculated as the absolute Michelson contrast between the luminance photon catches (sum input of the two LWS members of the double-cone) perceived by the prey species. Using light field measurements at 7 and 15 m depth, we reconstructed three scenarios in which scorpionfish and triplefin were likely to interact: (1) the scorpionfish and triplefin in the open against bright backgrounds where both PET and SAR eyeshine are present, (2) the scorpionfish in the open and the triplefin in the shade or against a dark background where only PET eyeshine is present (SAR eyeshine is weak or absent because of reduced side-welling illumination), and (3) the scorpionfish in the shade and the triplefin against a bright background in the open where only SAR eyeshine is induced (see Table S1 for details). The calculations generate values of chromatic contrast in just-noticeable-differences (JNDs), where values greater than one indicate discernible differences^30^, and values of achromatic contrasts as absolute Michelson contrasts (ranging from 0 to 1). To enhance camouflage, the contrast between the pupil and the iris should not only be reduced, but should also be comparable to the contrast of the overall patterning of the fish’s body. To assess the contrast between different skin patches on the body, we used the reflectance of four skin patches around the iris (calculated as the photon radiance of the skin normalized by the respective PTFE white standard facing the observer) measured in the field on the three scorpionfish. We calculated the absolute Michelson achromatic contrasts as perceived by the prey species (chromatic contrasts between the two pupils and the iris were not showing substantial differences) for all pairwise comparisons of the skin patches under the three scenarios at both depths. We then compared the achromatic contrasts between the irises and the natural and model pupils with the distribution of the skin patches achromatic contrasts. Visual models were implemented using the R package ‘pavo’ 1.0^32^.

### Units, statistics and data availability

Reflectance and transmittance are expressed as proportions, not percentages. Means are shown ± standard deviation unless specified otherwise. All data are available upon request.

## Results

### Occlusion of the *stratum argenteum* during dark-adaptation (laboratory)

Model validation didn’t show any problem. The reflectance of the light-adapted pupil of *Scorpaena porcus* was on average up to 30 times as strong as a white standard (29.7 ± 10.1). This value fell significantly during dark-adaptation to 5.1 *±* 0.4 (Fig. 2) with a log-linear regression coefficient of 0.0088 (95% CrI (Credible Interval): from 0.0065 to 0.0110). This implies that pupil reflectance decreased by 0.88% per minute. At the beginning of the experiment, one fish showed significantly higher reflectance, up to 40 times as strong as a white standard (95% CrI: from 0.3876 to 1.0685), but also a significantly faster decrease in reflectance of 1.25% per minute (95% CrI: from 0.0004 to 0.0069). The overall model fit was R^2^ = 0.84.

**Figure 2.**
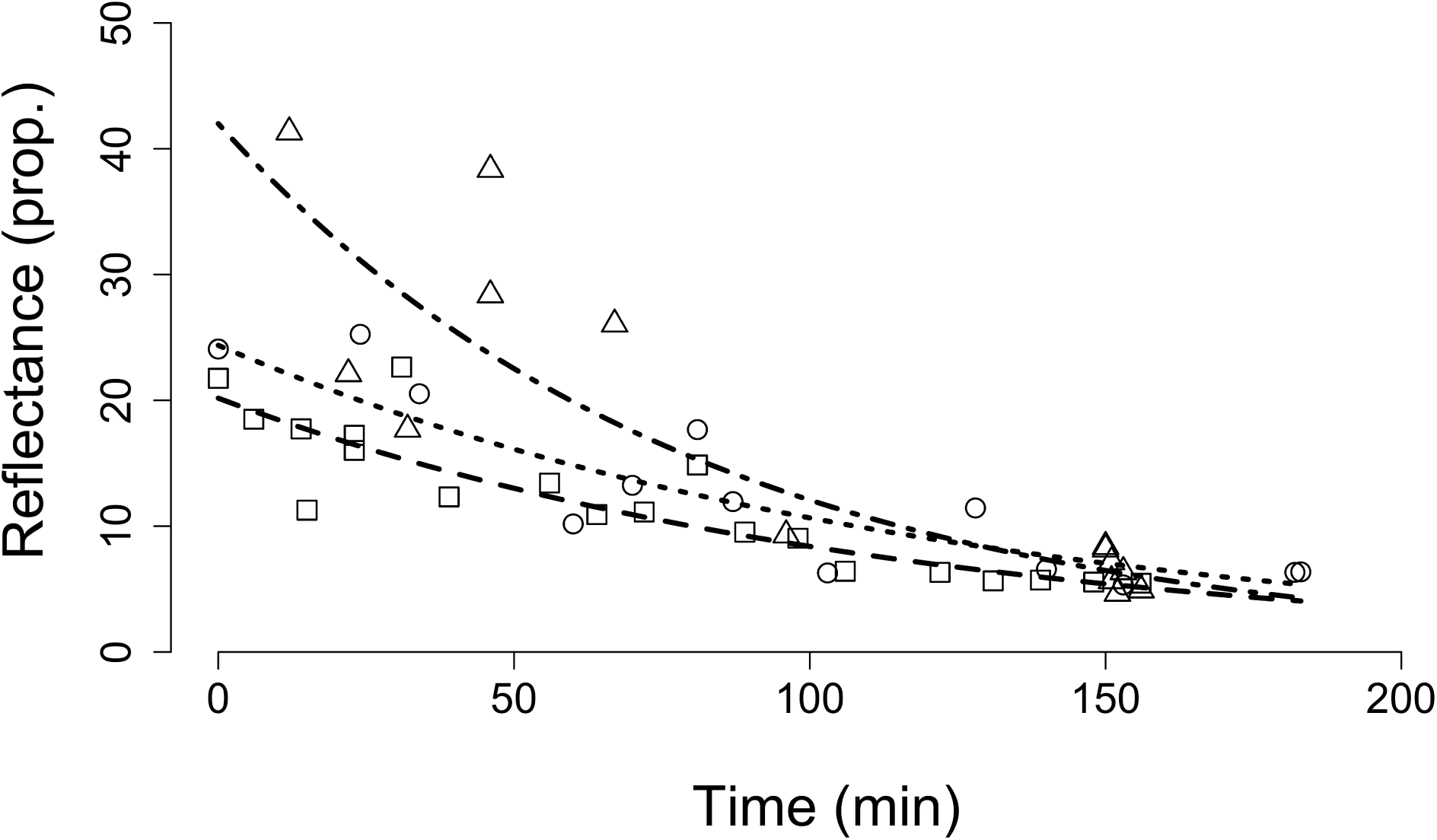
Reduction in eyeshine with increased dark-adaptation in *S. porcus.* SAR eyeshine total reflectance (pupil radiance normalized by white standard radiance) under coaxial illumination in the dark as function of time. At t = 0 the fish was light-adapted and the light was switched off resulting in total darkness. The small, coaxial source used to induce eyeshine was only switched on for a brief moment for each measurement. Radiance is integrated over 380-780 nm. Line styles and symbols indicate three different individuals.

### Contrast of natural and model pupil against iris (field)

The natural pupil showed reduced contrast against the iris relative to the model pupil, which was always considerably darker than the iris. At 7 m depth, this was true for both shaded and exposed conditions (Fig. 3). At 15 m depth, the difference between the pupils was strong only when exposed. Under shaded conditions both the natural and model pupil showed low contrast.

**Figure 3.**
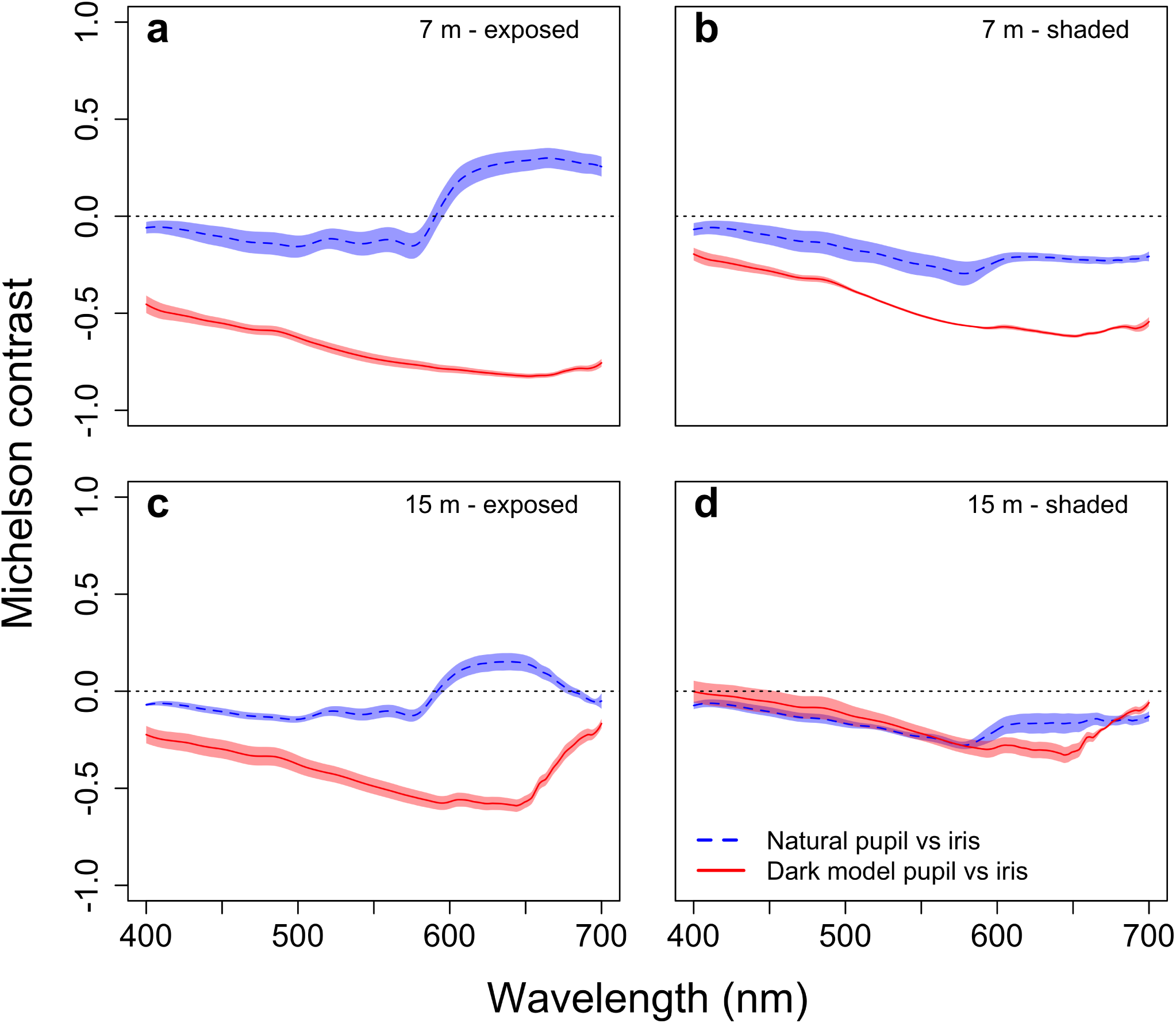
Michelson contrast at each wavelength between natural pupil or dark model pupil against the iris in the field. Measurements are from exposed and shaded *S. porcus* (n = 3 ind.) at 7 and 15 m depth. Positive values indicate that pupils are brighter. Shaded areas represent the standard error of the mean. **(a)** Exposed fish at 7 m. **(b)** Shaded fish at 7 m. **(c)** Exposed fish at 15 m. **(d)** Shaded fish at 15 m.

### PET and narrow-sense SAR eyeshine in light-adapted fish (laboratory)

Transmitted (PET) eyeshine (n = 10 ind.) had a transmittance of 0.033 *±* 0.028 (Fig. 4) with a shift towards the long wavelength part of the spectrum. Narrow-sense reflected (SAR) eyeshine (n = 10 ind.) was stronger than a PTFE white standard, showing a reflectance of 12.01 *±* 3.82 (Fig. 4).

**Figure 4.**
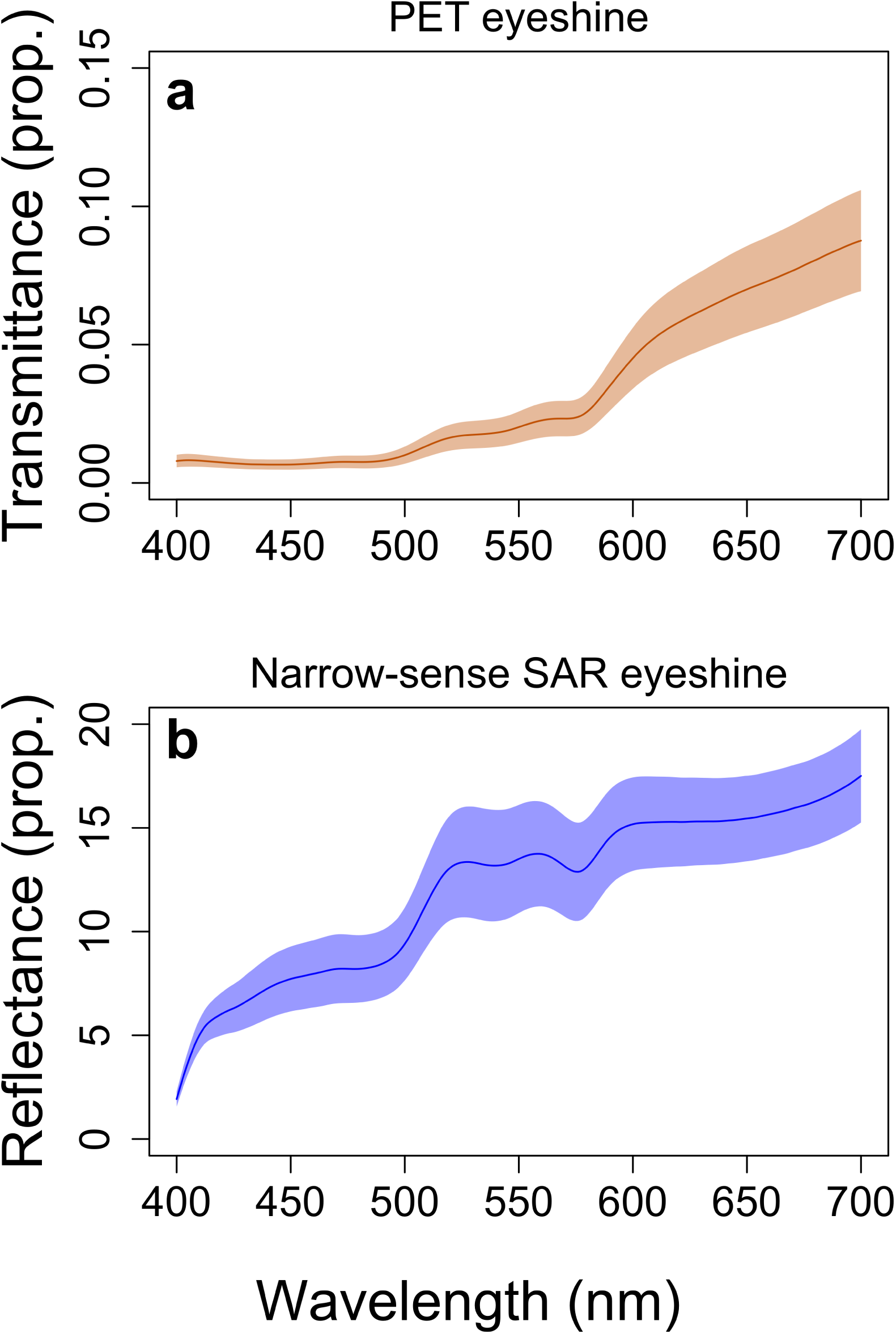
Contribution of two types of eyeshine in relation to wavelength in *S. porcus*. **(a)** PET eyeshine transmittance. **(b)** Narrow-sense SAR eyeshine reflectance. Y-values expressed as proportions of the pupil photon radiance normalized by the radiance of a PTFE white standard. Shaded areas represent the standard error of the mean.

### Broad-sense SAR eyeshine, iris and model pupil reflectance for visual modelling

Broad-sense SAR eyeshine (n = 9 ind.) yielded a reflectance of 0.06 *±* 0.01 (Fig. 5). The iris (n = 9 ind.) showed a reflectance of 0.15 *±* 0.05 (Fig. 5) and the model pupil a reflectance of the negligible value of 0.0008 *±* 0.0001 (not shown in the figure). Using these estimates (including PET transmittance), we predicted the pupil and iris radiance under natural light conditions and compared them to the actual field measurements (Fig.6). Overall, in fish exposed to ambient light, the averaged ratio between the measured and reconstructed values was 1.004 *±* 0.007 for the pupil and 1.012 *±* 0.012 for the iris, where a value of 1 represents the perfect match. When the fish was shaded, pupil and iris radiances were predicted with a ratio of 1.024 *±* 0.009 and 1.008 *±* 0.008.

**Figure 5.**
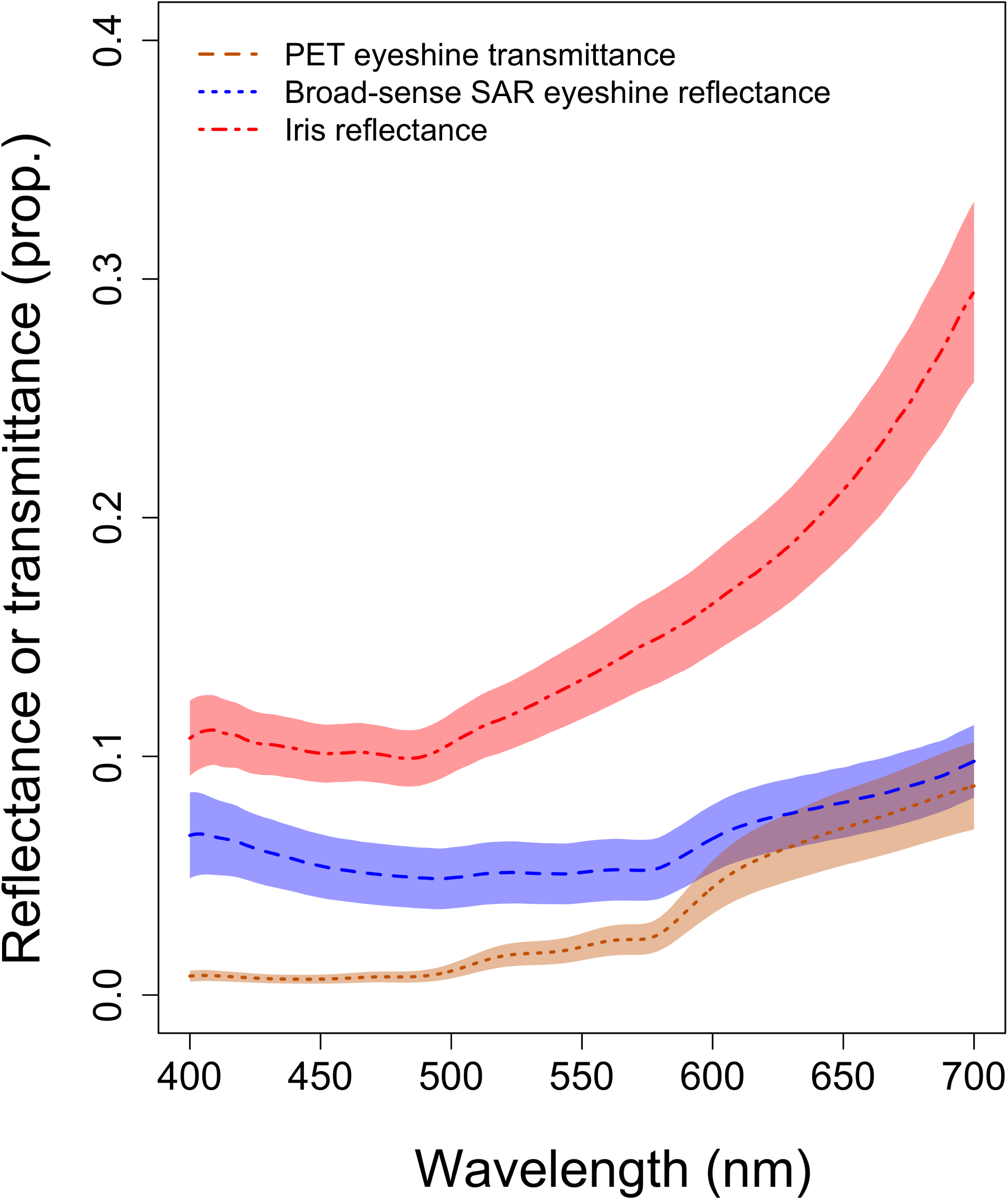
Laboratory estimates to inform visual models. Broad-sense SAR eyeshine reflectance, PET eyeshine transmittance (as in Fig. 4) and iris reflectance. Parameters are normalized by the associated white standard measurements. Shaded areas represent the standard error of the mean.

**Figure 6.**
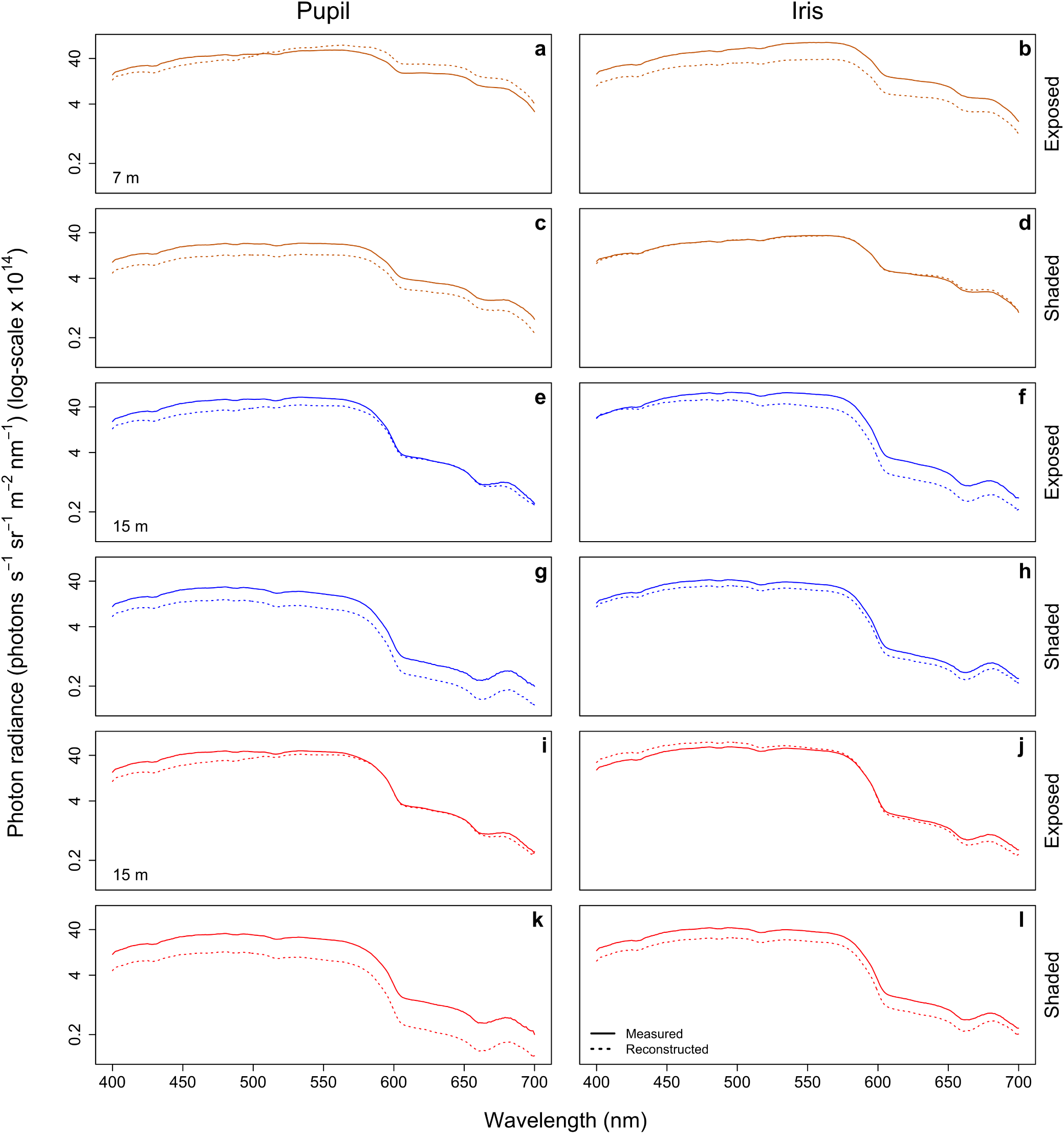
Reconstruction of known field pupil and iris radiance. Photon radiance of **(a,e,i)** pupil and **(b,f,j)** iris in exposed *S. porcus* individuals. Panels **(c,g,k)** and **(d,h,l)** show the same for shaded individuals. Solid lines show radiance measured in the field. Dotted lines show radiance reconstructed using the ambient light measurements for each fish combined with the eyeshine and iris relative radiances estimated in the laboratory. Y-values are expressed on a log-scale. Each colour indicates a different individual, placed at 7 or 15 m depth.

### Visual models to assess pupil camouflage as perceived by prey fish

A natural pupil with PET and/or SAR eyeshine showed a similar chromatic contrast against the iris if compared to a model pupil in all scenarios, except for a slight increase when only PET eyeshine was present (triplefin shaded, scorpionfish exposed) (Tab. 1). However, achromatic contrast was substantially reduced (Tab. 1). Whereas the model pupil without eyeshine against the iris showed absolute achromatic contrasts around 1, the contribution of one or both eyeshine mechanisms reduced this contrast to less than 0.4. This value is well within the range (from 0 to 0.6) of the achromatic contrasts between skin patches found near the irises (Fig. 7).

**Table 1.**
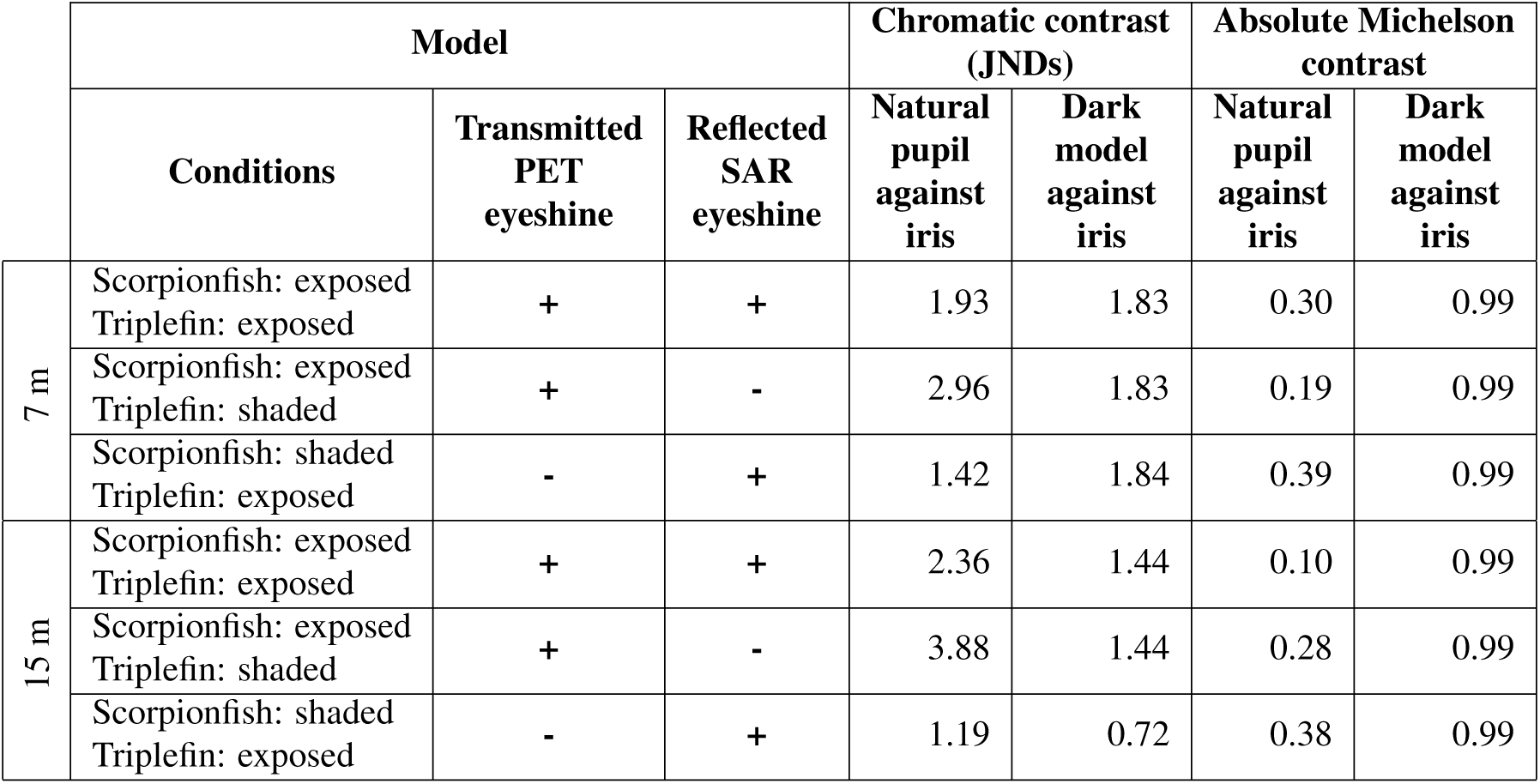
Estimated chromatic and achromatic contrast in the eye of a scorpionfish. Chromatic contrast values in just-noticeable-differences are calculated between the natural pupil (with eyeshine), the model pupil (without eyeshine) and the iris for three light scenarios (described by cols. 1-3) at two depths. Achromatic contrast values are expressed as absolute Michelson contrasts between the same three structures. All contrasts are calculated from the perspective of the triplefin *Tripterygion delaisi*, a common prey species. SAR eyeshine refers to “broad-sense” SAR eyeshine. See Material and Methods and Table S1 for details.

**Figure 7.**
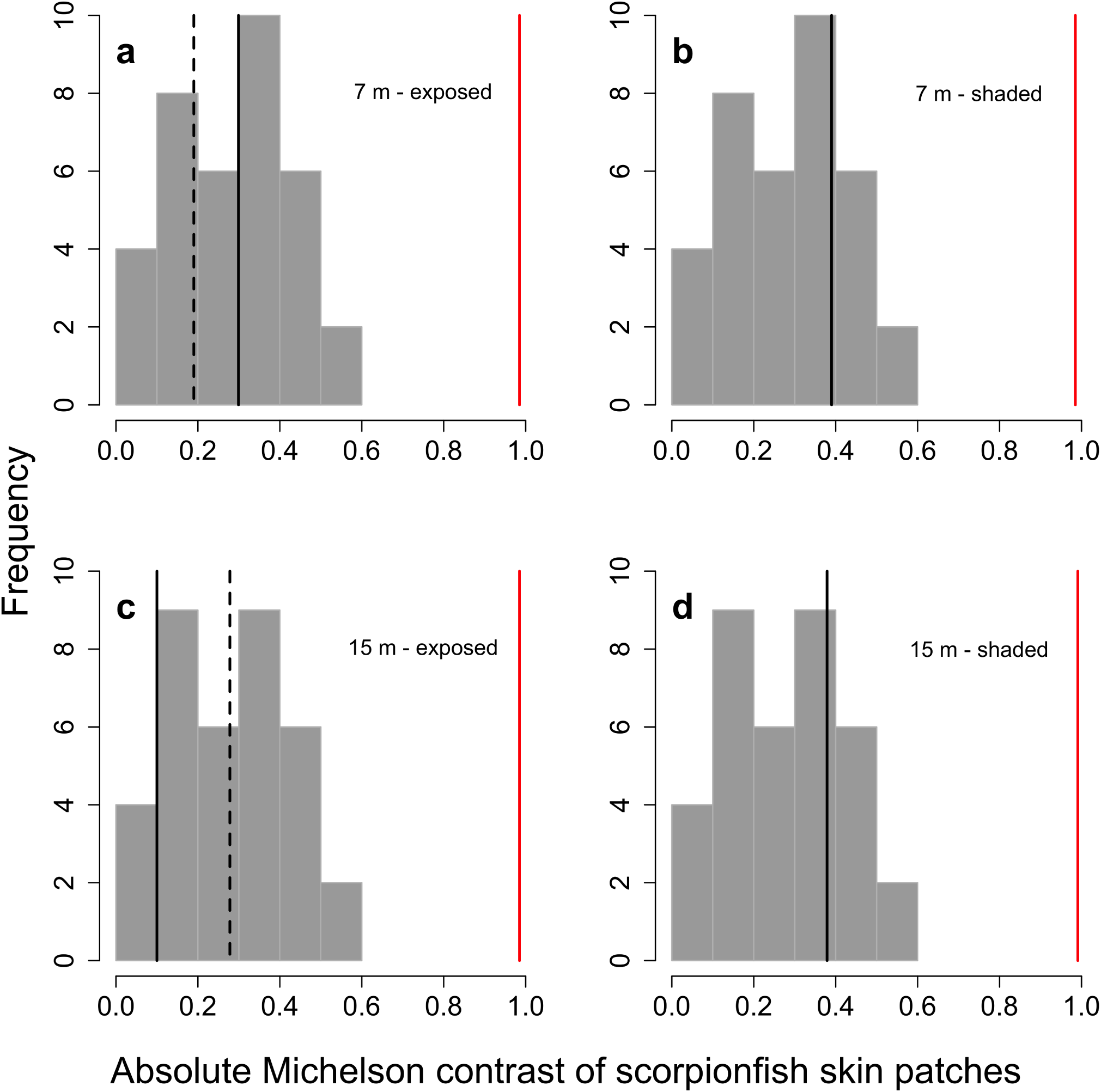
Predicted absolute achromatic contrast of the pupil with and without eyeshine against the iris of *S. porcus* compared to the contrast among body patches as perceived by a prey species. The achromatic contrast between a natural pupil with eyeshine and its iris (black lines) was small and fell within the distribution of the achromatic contrasts between skin patches (all combinations of four patches in each of the three fish measured). This was not the case when eyeshine was prevented, as in the dark model pupil (red lines). Black lines indicate contrast values between the natural pupil and the iris when *T. delaisi* is shaded (dashed) or exposed (solid) for four *S. porcus* scenarios: **(a)** exposed and **(b)** shaded at 7 m depth, **(c)** exposed and **(d)** shaded at 15 m depth. Red lines indicate contrast values between the black model pupil and the iris when *T. delaisi* is shaded (dashed) or exposed (solid) for the same four *S. porcus* scenarios (note that in scenario (a) and (c) two lines overlap).

## Discussion

Only few families of cryptobenthic fish occlude their ocular reflector in the dark and expose it during the day, which is opposite to what is known from most organisms. Reversed occlusion can make a pupil bright during the day, but keeps it dark at night, even when illuminated. We confirmed the presence of reversed occlusion in the scorpionfish *Scorpaena porcus* and show that two different eyeshine types contribute to daytime pupil radiance: *Stratum Argenteum Reflected* (SAR) and *Pigment Epithelium Transmitted* (PET) eyeshine. Spectroradiometry in the field shows that eyeshine might reduce the contrast between pupil and iris, which may help to conceal an otherwise black pupil during the day. Visual modelling confirmed this observation: daytime eyeshine reduces pupil contrast against the surrounding tissue, decreasing the detectability of *S. porcus* pupil by the perspective a prey species under three light scenarios where SAR and PET eyeshine differ in their contribution to pupil radiance. The presence of an occluded *stratum argenteum* in the dark-adapted eye was first observed in *S. porcus* during night dives (pers. obs.) and is in line with published observations on toadfishes^9^. Why these fishes suppress eyeshine when dark-adapted remains unclear. Using the *stratum argenteum* as a reflector in dim light conditions would allow for an increased quantal catch and thus increased visual acuity^9^. Covering it with pigmentation, however, forgoes this option. A possible explanation could be that a dark pupil under low light is just one end of a fine-tuning scale for pupil brightness under changing light conditions in the course of the diurnal cycle. Future studies where eyeshine is measured under a range of light intensities could elucidate whether this is the case. Enhanced pupil camouflage by reversed occlusion may come at the expense of reduced visual acuity due to a possible increase in internal scatter in the light-adapted eye and a reduced photon catch in the dark-adapted eye. On the other side, however, enhanced pupil camouflaged by means of eyeshine may explain why scorpionfish possess unobstructed, large pupils. Most other cryptobenthic predators feature fringes, skin flaps, or have small eyes to reduce their eyes conspicuousness.

Daytime eyeshine due to the absence of melanocytes in the choroid has been already described for toadfishes and scorpionfishes^6,9,13^. Until now, however, the focus was exclusively on reflected (SAR) eyeshine^6,9,13^. Our measurements in the laboratory show that eyeshine in *S. porcus* has a second component in the form of transmitted (PET) eyeshine. These two eyeshines could be mechanistically linked because they both rely on the translucent choroid and the exposed *stratum argenteum*. Since the combination of PET and SAR eyeshine could enhance pupil camouflage, the bright pupils of this fish should not be seen just as a side-effect of eye anatomy, but rather as a possible adaptive trait.

For a sit-and-wait predator, successful prey capture strongly depends on its crypsis^33^. In this study, we show that daytime eyeshine enhances the camouflage of the pupil in a cryptobenthic predator. Eyeshine does not perfectly camouflage the eye, but it likely reduces the probability of detection compared to a situation in which the pupil is dark. The contribution of SAR eyeshine to pupil brightness had already been summarily described^6^, but we also showed that transmitted PET eyeshine is an additional component that makes a significant contribution, regardless of its low transmittance, particularly because it is induced by the strong down-welling light rather than the weaker side-welling light. A similar concealing process has been described for pelagic stomatopod crustacean larvae that use a photonic structure external to the optical pathway of the eye to hide their dark retinas^23^. Since in *S. porcus* the *stratum argenteum* is located in the optical pathway, this ocular reflector may also improve visual sensitivity in dim light environments^9^. However, reversed occlusion of the *stratum argenteum* strongly suggests that this function is secondary to camouflage.

We did not inform our visual models with the true physical properties of (narrow-sense) SAR eyeshine. This parameter showed extremely high reflectance values (Fig. 4), as expected in the presence of a strong reflector. However, it can only be included if the observer possesses a light source close to its eyes in an otherwise dark environment. Under these conditions, the target pupil would appear much brighter and might stand out against the iris, potentially decreasing eye camouflage. This possibility is to be addressed in future work. Our models instead used the broad-sense SAR eyeshine reflectance, which represents the more general situation in which the complete side-welling light field is considered and SAR eyeshine is not actively induced.

This is the first study showing that a vertebrate can produce daytime eyeshine by means of two complementary mechanisms to reduce the conspicuousness of its large pupils to one of its prey species. We propose that the unusual reversed occlusion of its ocular reflector has evolved in response to selection to optimize the trade-off between camouflage and vision.

## Acknowledgements

We warmly thank the STARESO staff for hosting and supporting our research group in their field station, Oeli Oelkrug for fish maintenance at the University of Tuebingen and Nils Anthes for useful comments on a previous draft.

## Author contributions

MS conceptualised the study. MS and NKM designed the experimental setups. MS performed spectroradiometry in the laboratory. MS, UKH and NKM performed spectroradiometry in the field. MS analysed the data. MS and P-PB implemented the visual models. MS drafted the manuscript. MS, NKM and P-PB edited the manuscript and all authors commented on the completed manuscript.

